# The pro-inflammatory cytokines IFN-α and TNF-α inhibit organoid-derived extravillous trophoblast invasion

**DOI:** 10.1101/2025.08.15.670497

**Authors:** A. Jantine van Voorden, Fangxu Lin, Souad Boussata, Remco Keijser, Liana Barenbrug, Bente Horselenberg, Ans M. M. van Pelt, Wendy Dankers, Susana M. Chuva de Sousa Lopes, Gijs B. Afink

## Abstract

Proper placental development requires differentiation and invasion of extravillous trophoblasts (EVTs) into the maternal decidua to ensure adequate remodeling of spiral arteries and support fetal growth. Perturbations in these processes are associated with pregnancy complications such as preeclampsia and fetal growth restriction. The incidence of these pregnancy complications is increased in women with immune-mediated inflammatory diseases, which are characterized by elevated levels of pro-inflammatory cytokines, such as interferon alpha (IFN-α) and tumor necrosis factor alpha (TNF-α). However, the direct effects of these cytokines on trophoblast differentiation and invasion remain unclear. Using human trophoblast organoid models, we demonstrate that IFN-α and TNF-α impair EVT invasion while preserving their differentiation capacity. High-resolution imaging of untreated organoid–decidua co-cultures revealed extensive invasion of organoid-derived trophoblasts into the decidual stroma and spiral arteries. In contrast, organoids treated with IFN-α, in particular, exhibited markedly reduced invasion within this system. Transcriptome profiling indicated altered expression of several pathways involved in invasion upon cytokine treatment. These findings support the notion that elevated levels of IFN-α and TNF-α can directly impair trophoblast invasive capacity, potentially contributing to suboptimal placental development and adverse pregnancy outcomes in the context of inflammatory disorders.

## Introduction

Severe pregnancy complications, including early-onset preeclampsia and fetal growth restriction, are primarily caused by defective placenta development (1, 2). The placenta provides the fetus with oxygen and nutrients, and eliminates waste products. Maternal-fetal exchange is mediated by syncytiotrophoblasts, multinucleated cells derived from fusion of cytotrophoblasts, which are in direct contact with the maternal blood. At the tips of anchoring placental villi, cytotrophoblasts differentiate into extravillous trophoblasts (EVTs), a process that involves epithelial–mesenchymal transition (EMT), and results in an invasive cell type (1–5). Subsequently, EVTs infiltrate the maternal decidua, and actively remodel the spiral arteries, converting them into high-conductance, low-resistance blood vessels. Spiral artery remodeling is characterized by loss of smooth muscle cells and replacement of endothelial cells by EVTs, and is regulated by EVTs as well as decidual natural killer cells (3, 6). EVTs can reach the spiral arteries by migration through the decidual stroma (interstitial EVTs) or migration directly from the anchoring villi into the arteries (endovascular EVTs). The processes of EVT differentiation, invasion and spiral artery remodeling are essential for facilitating an adequate uteroplacental blood flow, and proper maternal-fetal exchange; a dysregulation in these processes can result in adverse pregnancy outcomes (1–3, 7–9).

The maternal immune system plays an important role throughout pregnancy, and needs to adapt to the developing embryo and later fetus (10, 11). During the peri-implantation period, a pro-inflammatory microenvironment (Th1/Th17 response) is required to allow blastocyst implantation. However, to avoid rejection of the semi-allogeneic fetus, an anti-inflammatory microenvironment (Th2/regulatory T cell response) becomes dominant during further fetal development. At the end of pregnancy, a pro-inflammatory microenvironment is again established to prepare for parturition. These shifts are tightly regulated by immune cells through the secretion of both pro-inflammatory and anti-inflammatory cytokines. A disturbed immune balance is associated with pregnancy complications, in particular preeclampsia, which is accompanied by systemic inflammation and endothelial cell dysfunction (10–14).

Chronic diseases in which the maternal immune system is inherently dysregulated, such as immune-mediated inflammatory diseases, pose an increased risk for adverse pregnancy outcomes (15–17). These diseases are a clinically heterogeneous group of conditions typically characterized by chronic inflammation and organ damage, which are caused or accompanied by overactivity of pro-inflammatory cytokines (18, 19). Immune-mediated inflammatory diseases especially affect women, many of whom are of reproductive age, emphasizing the relevance of their effect on pregnancy outcomes. However, it remains unclear whether the involved cytokines directly affect early placenta development, and whether that contributes to the increased risk for pregnancy complications.

Systemic lupus erythematosus (SLE) is an immune-mediated inflammatory disease that is characterized by high activity of type I interferons (IFNs), in particular IFN-α (20, 21). Women with SLE have an increased risk for adverse pregnancy outcomes, including preeclampsia, preterm birth and low birth weight (16, 22). In addition, SLE pregnancies are associated with lower placental weight and abnormalities in placental vascularity (23). Moreover, increased IFN-α activity correlates with the onset of preeclampsia in SLE patients (22–24), and elevated IFN-α plasma levels have been associated with lower birth weight (25). As IFNs play an important role in the response to viral and bacterial infections, they are crucial for protection of the fetus from pathogens in normal pregnancies (26). However, the effect of increased IFN-α levels on early placentation events has been poorly studied. IFN-β, another type I IFN, was shown to inhibit fusion of trophoblasts into syncytiotrophoblast in a study using the BeWo trophoblast cell line (27). In addition, a recent publication showed that IFN-β inhibits EVT invasion by targeting EMT, in an organ-on-a-chip model using primary trophoblasts and endothelial cells (28). As IFN-α and IFN-β bind to the same receptor, it is plausible that IFN-α has a similar effect on trophoblasts.

Tumor necrosis factor alpha (TNF-α) is a pro-inflammatory cytokine that is highly secreted in several immune-mediated inflammatory diseases, including SLE, rheumatoid arthritis, psoriasis and inflammatory bowel disease (29, 30). Aberrant levels of TNF-α during pregnancy are associated with preeclampsia and other adverse pregnancy outcomes (31). In pregnant mice, TNF-α infusion induces hypertension and proteinuria, resembling preeclampsia (32). Several studies using first-trimester placental explants, primary EVTs or trophoblast cell lines, have suggested that TNF-α inhibits EVT invasion, by targeting processes such as extracellular matrix degradation or adhesion, apoptosis, proliferation and/or differentiation (33–40). However, the exact mechanisms remain unclear.

Our possibilities to study early placenta development in vitro have increased since the derivation of human trophoblast stem cells (TSCs) from first-trimester placental tissue (41) and the generation of trophoblast organoids (42, 43). In contrast to other model systems, such as choriocarcinoma cell lines and primary cells, TSCs have the capacity of long-term self-renewal and differentiation into hormone-producing syncytiotrophoblasts and invasive EVTs (41). Moreover, they fulfill the additional criteria for bona fide TSCs as proposed by Lee *et al*. (44) and Karvas *et al*. (45). By seeding TSCs into an extracellular matrix, they self-organize into three-dimensional structures, consisting of a cytotrophoblast layer partly filled with syncytiotrophoblasts, and can be induced to give rise to EVTs (46). Because of their invasive nature, the EVTs invade the extracellular matrix, forming protrusions of cells with mesenchymal morphology. Trophoblast organoids therefore serve as a valuable system to study EVT invasion in vitro.

In this study, we investigated in detail the impact of IFN-α and TNF-α on EVT differentiation and invasion using monolayer TSC cultures, trophoblast organoids, and first-trimester placental and decidual tissue. Our findings indicate that IFN-α and TNF-α do not interfere with EVT differentiation, but significantly suppress EVT invasion into both Matrigel matrices and maternal decidual tissue. For the latter, we co-cultured trophoblast organoids with decidual explants, in which we observed pronounced trophoblast invasion into the stroma and arteries using high-resolution optical sectioning. Transcriptome analysis on Matrigel-invading EVTs suggested that EMT and other pathways associated with cellular invasion are affected by these cytokines, but direct targets that interfere with these pathways remain to be elucidated. As EVT invasion is crucial for proper spiral artery remodeling and a healthy pregnancy, the direct effect of IFN-α and TNF-α on trophoblasts may contribute to the increased risk for adverse pregnancy outcomes in patients with elevated IFN-α/TNF-α activity.

## Results

### IFN-α and TNF-α do not affect monolayer EVT differentiation

For this study, we made use of a TSC line previously generated from first-trimester placental tissue (46). This TSC line, as well as additionally generated TSC lines, express the genes encoding IFN-α receptors (*IFNAR1* and *IFNAR2*) and TNF-α receptors (*TNFRSF1A* and *TNFRSF1B*) (47). While *TNF* mRNA was endogenously expressed at very low levels in TSCs, transcripts encoding IFN-α were not detected.

To assess the impact of the cytokines on monolayer EVT differentiation, IFN-α or TNF-α was added to the EVT differentiation medium throughout the entire culture period. Irrespective of these treatments, the cells obtained an elongated, mesenchymal morphology typical for EVTs (Fig. 1A). Moreover, the cells exhibited clear downregulation of TSC markers and upregulation of EVT markers relative to undifferentiated TSCs (Fig. 1B), suggesting that IFN-α and TNF-α do not block EVT differentiation.

**Fig. 1.**
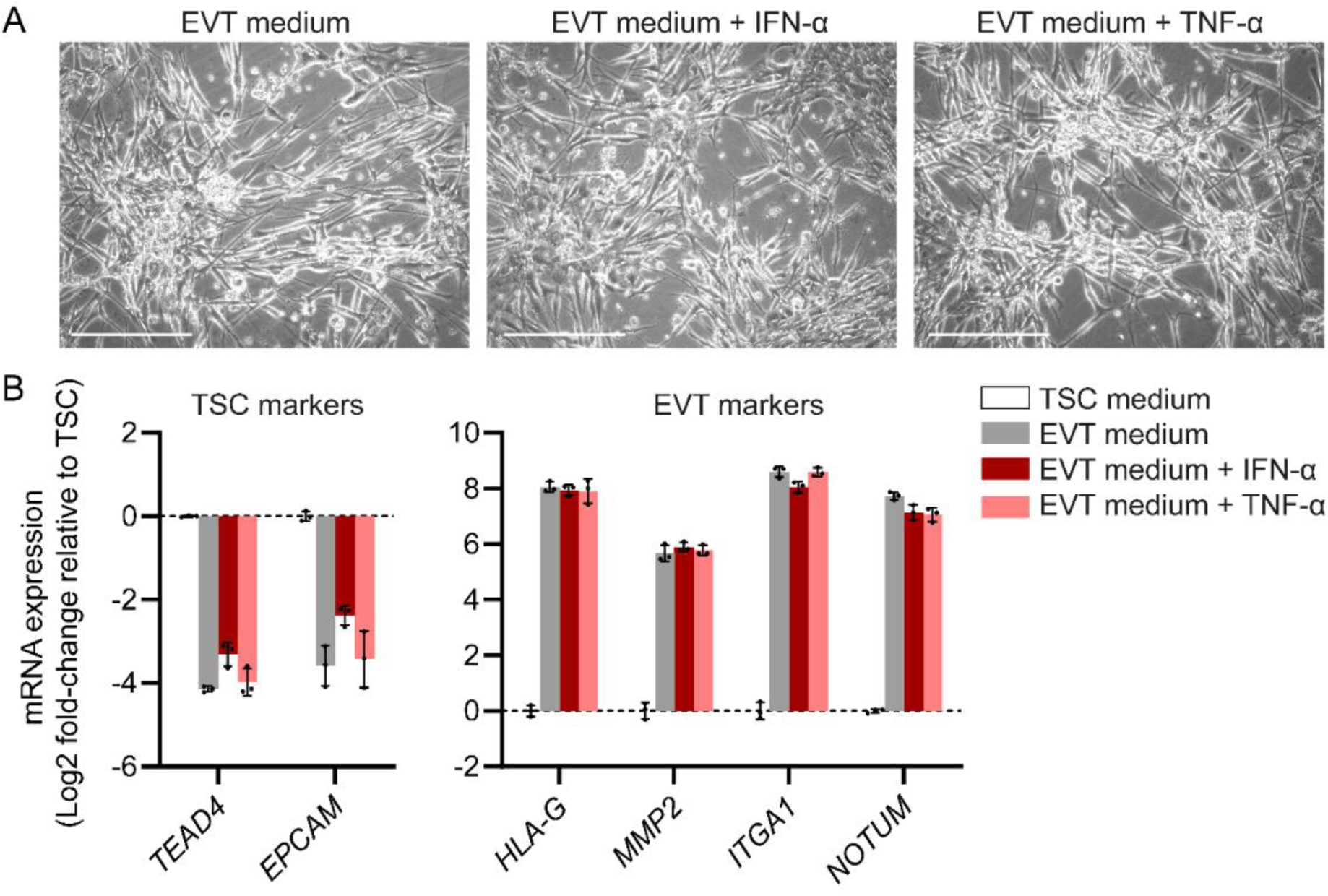
IFN-α and TNF-α do not hinder monolayer EVT differentiation. TSCs were induced to undergo monolayer EVT differentiation for 8 days in the presence of IFN-α (5 ng/ml) or TNF-α (10 ng/ml), or without treatment as a control. (A) Representative phase-contrast images at EVT differentiation day 8. Scale bars: 400 µm. (B) TSC and EVT marker mRNA expression measured by RT-qPCR. Bars represent mean log2 fold-change ± SD relative to undifferentiated TSCs (n=3 independent experiments). TSC markers were significantly downregulated and EVT markers significantly upregulated *(P* < 0.01) in EVT conditions relative to the TSC condition. Differences between treatments were not significant.

### IFN-α and TNF-α inhibit the invasive capacity of EVTs into Matrigel matrix

Similarly, we added the cytokines to trophoblast organoid medium (TOM), overlaying TSCs seeded in Matrigel, to investigate whether TSCs are able to form organoids under these conditions. Morphology of the organoids seemed normal, and they showed similar mRNA expression of TSC, EVT and STB markers as untreated organoids (Fig. S1A-B). Thus, IFN-α and TNF-α did not seem to affect trophoblast organoid formation.

However, when adding the cytokines to organoids in trophoblast organoid (TO)-EVT medium to investigate the effect on EVT invasion, we observed reduced EVT outgrowth from the organoids into the Matrigel matrix (Fig. 2A). This was not attributable to impaired EVT differentiation, as mRNA levels of EVT markers were markedly elevated compared with organoids in TOM, and similar to those of organoids in TO-EVT medium without cytokines (Fig. 2B). Furthermore, immunostaining for HLA-G and MMP2 confirmed the presence of EVTs; however, these cells remained in closer proximity to the organoid bodies (Fig. 2A, Fig. S2), indicating a reduced invasion capacity into the surrounding Matrigel matrix in the presence of IFN-α or TNF-α.

**Fig. 2.**
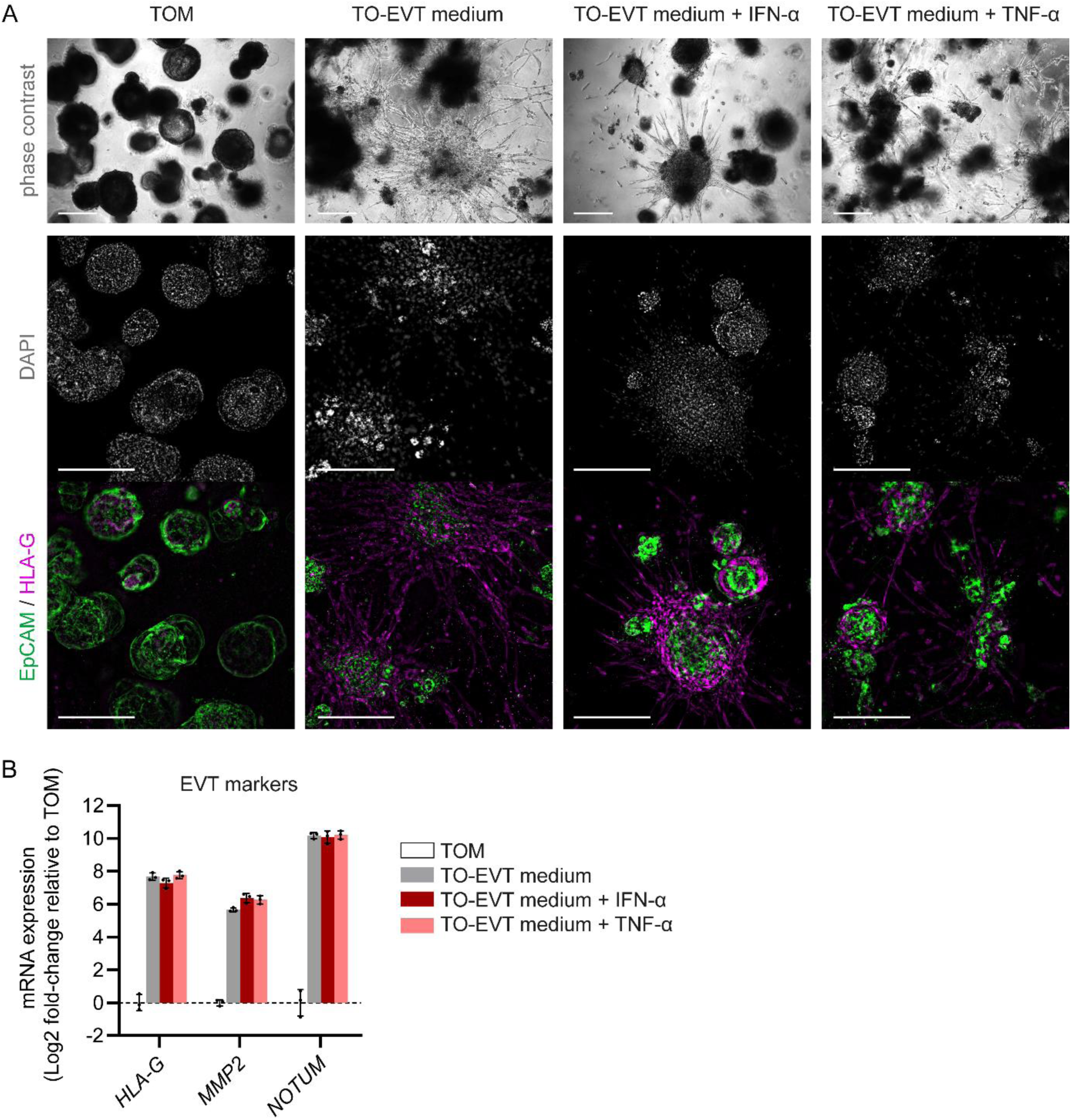
IFN-α and TNF-α inhibit EVT invasion but do not hinder EVT differentiation in trophoblast organoids. Organoids were grown in Matrigel domes in trophoblast organoid medium (TOM) for 5 days, and consecutively induced to undergo EVT differentiation in trophoblast-organoid(TO)-EVT medium in the presence or absence of IFN-α or TNF-α for 14 days, or retained in unsupplemented TOM. (A) Representative phase-contrast images and immunofluorescence extended depth of field composite images. Organoids were stained for EpCAM (TSC marker, green), HLA-G (EVT marker, magenta) and DAPI (nuclei, white). Scale bars: 400 µm. Additional immunofluorescence images are shown in Fig. S2. (B) EVT marker mRNA expression measured by RT-qPCR. Bars represent mean log2 fold-change ± SD relative to untreated organoids in TOM (n=3 independent experiments). For all EVT conditions, EVT marker expression was significantly increased *(P* < 0.001) compared with the TOM condition.

Using placental villous explants, we observed a similar effect: untreated explants developed prominent outgrowth composed of elongated EVTs, whereas explants treated with IFN-α or TNF-α exhibited altered outgrowth patterns characterized by small, rounded EVTs closely associated with the villous structures (Fig. 3, Fig. S3).

**Fig. 3.**
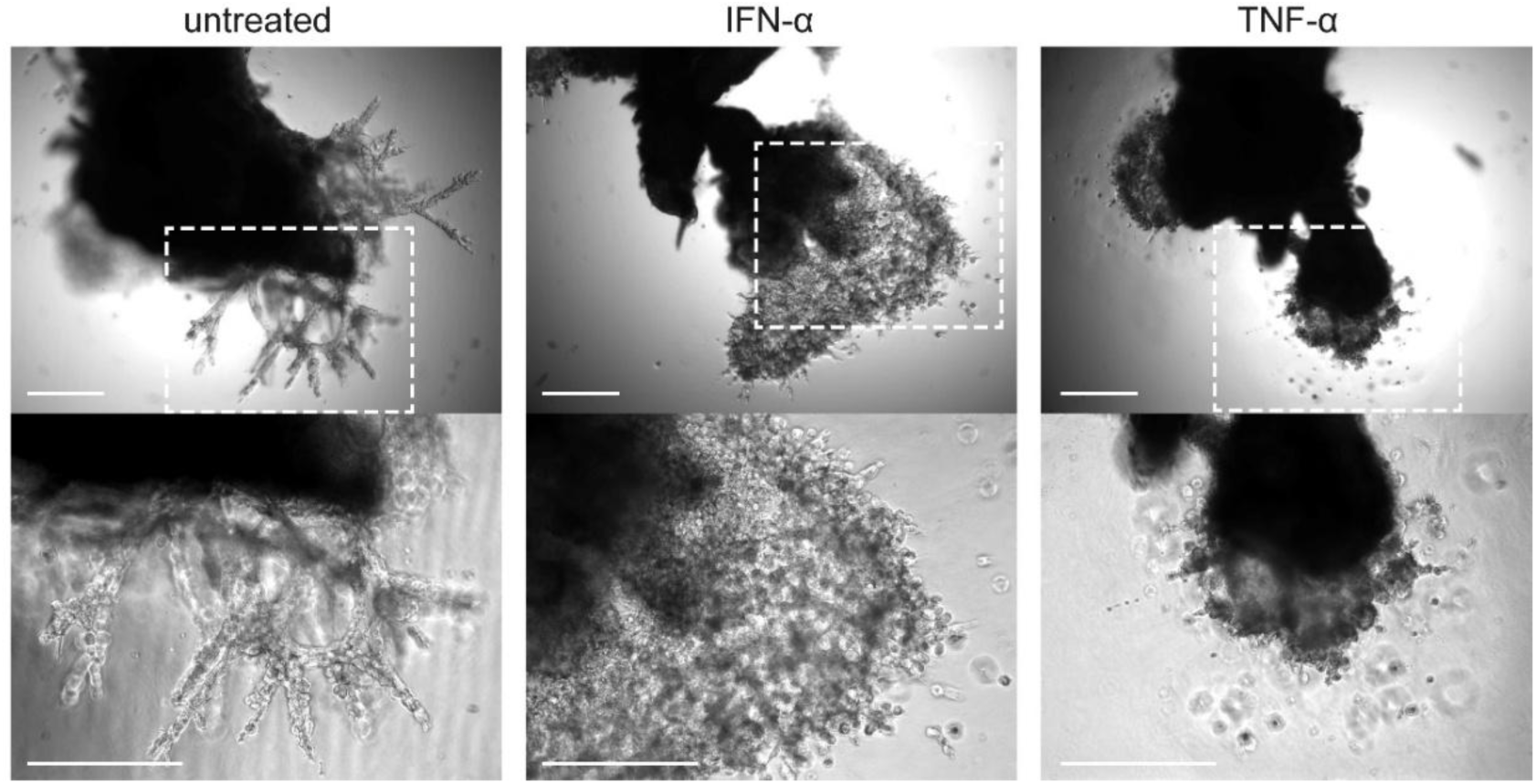
IFN-α and TNF-α affect EVT invasion in first-trimester placental villous explants. Representative phase-contrast images of placental villous explants (8 wks gestation), which were seeded on Matrigel and treated with IFN-α or TNF-α. Higher magnification images of the indicated areas are shown in the bottom row. Scale bars: 400 µm. Representative images obtained from other tissue donors are shown in Fig. S3.

To be able to quantify the invasive capacity of EVTs, we set up a single-organoid system. In this, we induced organoid formation and early EVT differentiation within Matrigel domes for 5 days each, and subsequently transferred the organoids individually onto a Matrigel layer in separate wells, in which they were induced to undergo EVT differentiation for another 9 days. In this system, EVTs invaded the Matrigel as rounded, individual cells, and staining with the live cell marker calcein AM showed that the majority of these cells were alive (Fig. 4A). Measuring the distance of invading EVTs to the surface of the organoid body demonstrated that treatment with IFN-α or TNF-α indeed reduced EVT invasion capacity (Fig. 4B). Additionally, the number of invasive EVTs, i.e. EVTs detached from the organoid body, was significantly reduced (Fig. 4C).

**Fig. 4.**
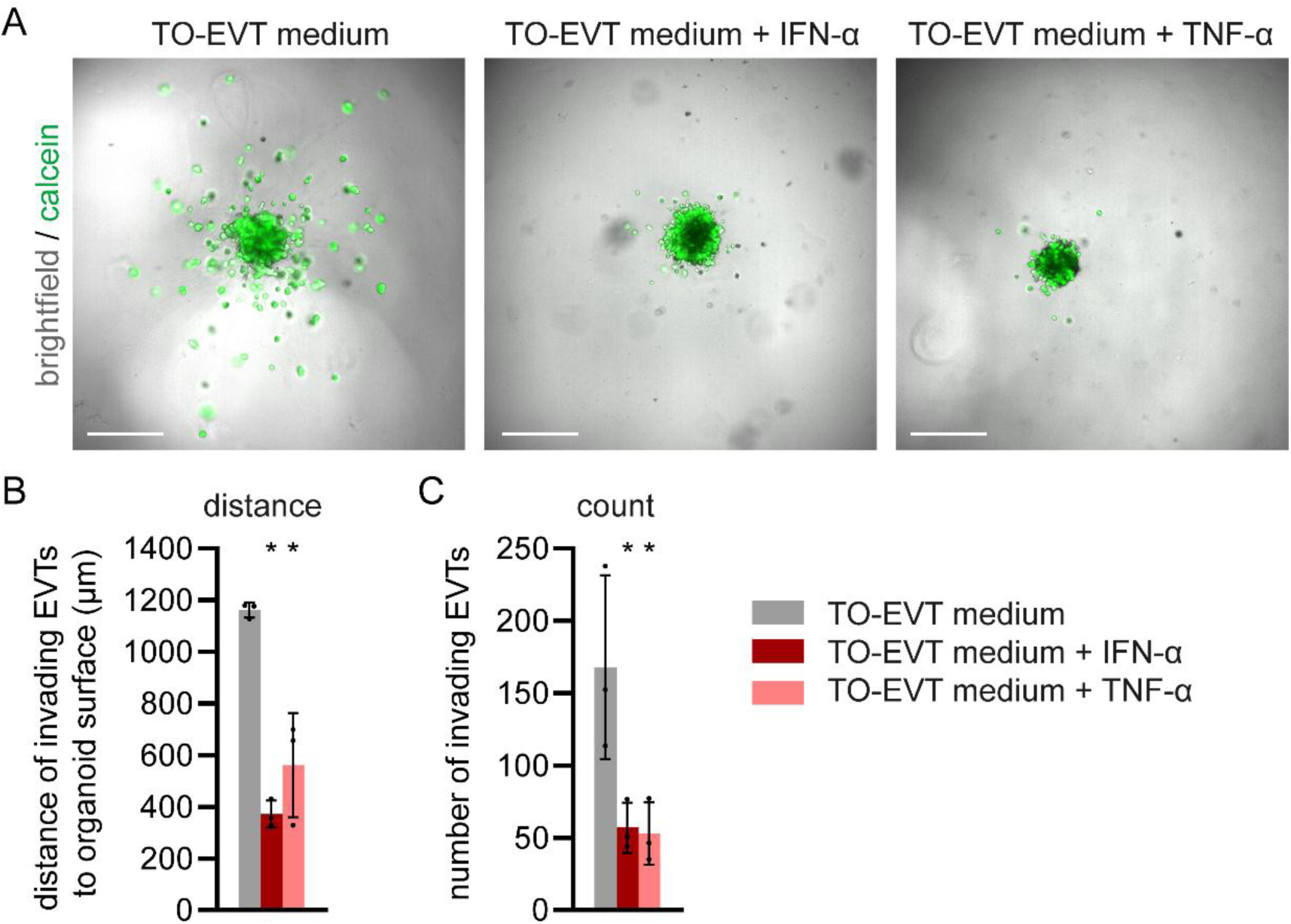
Quantification of EVT invasion using single organoids demonstrates that IFN-α and TNF-α inhibit EVT invasion. Trophoblast organoids were transferred individually onto separate wells on top of a layer of Matrigel in TO-EVT medium, with or without IFN-α/TNF-α (10 organoids per condition per experiment). Single organoids were stained with calcein AM, imaged, and analyzed in three dimensions. (A) Representative extended depth of field composite images of Z stacks displaying calcein (green) and brightfield. Scale bars: 500 µm. (B) Mean distance of invading EVTs (using mean of 5 furthest migrated EVTs per image) to surface of organoid body ± SD. (C) Mean number of invading EVTs ± SD. Data points represent n=3 independent experiments. \**P* < 0.001, compared with untreated control.

### IFN-α and potentially TNF-α inhibit EVT invasion into decidua

To investigate whether IFN-α and TNF-α also disrupt EVT invasion in the context of the maternal decidua, we established a co-culture system of trophoblast organoids with ex vivo first-trimester decidual parietalis tissue. The organoids were generated from TSCs expressing green fluorescent protein (GFP) under control of a cytomegalovirus (CMV) promoter, resulting in constitutive GFP reporter expression in all cell types. This allowed observation of trophoblast invasion into the decidual tissue. Prior to co-culture, the organoids were cultured for ∼1 week in TO-EVT medium to induce EVT differentiation, with or without addition of the cytokines. Small fragments of decidua parietalis were transferred onto a layer of Matrigel, and overlayed with the organoids suspended in TO-EVT medium, with or without added cytokines (Fig. 5A). Co-cultures were incubated for 8-11 days.

**Fig. 5.**
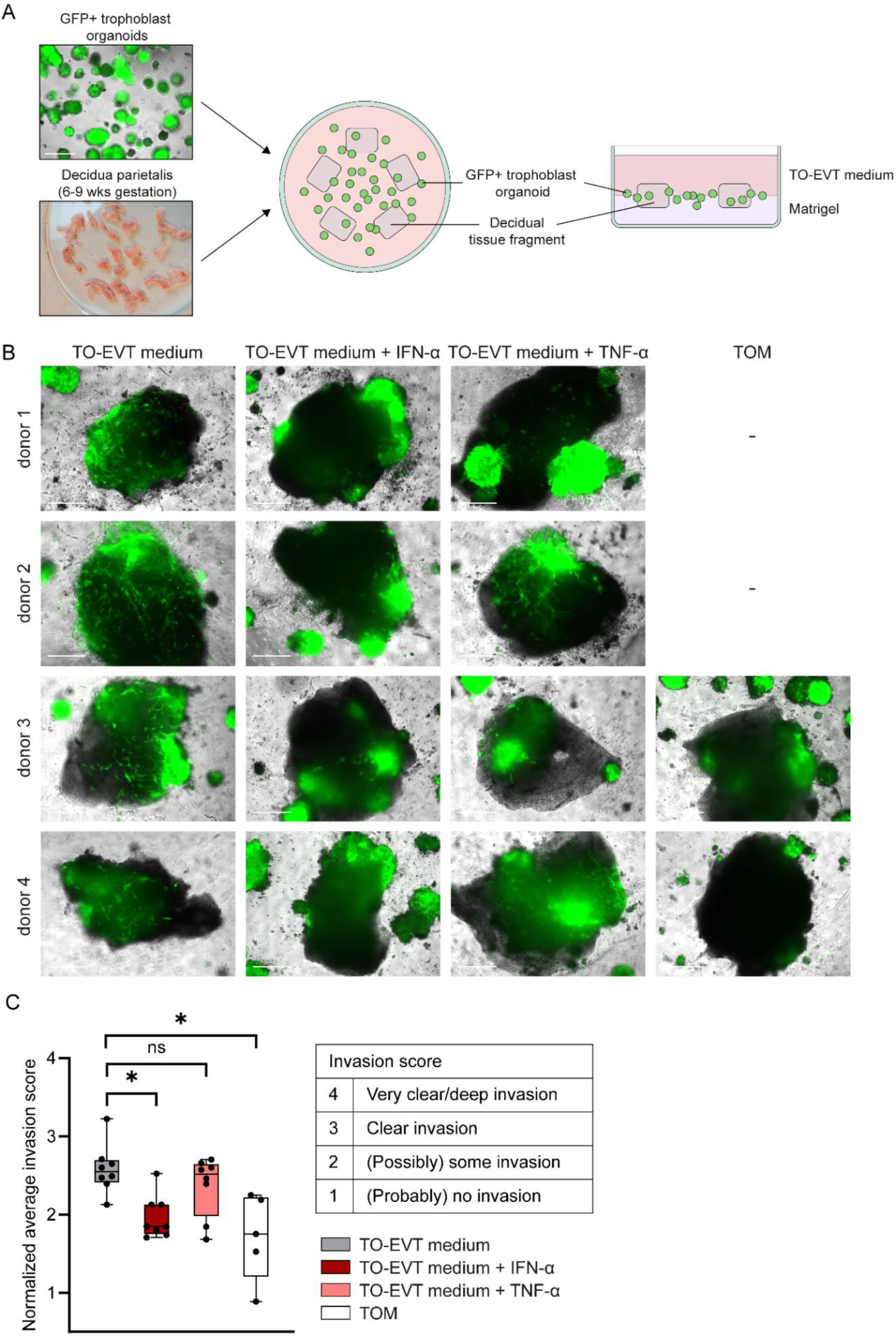
IFN-α and potentially TNF-α inhibit EVT invasion from trophoblast organoids into first-trimester decidual tissue. (A) Schematic drawing of the experimental set-up. GFP-positive trophoblast organoids were co-cultured with decidual parietalis tissue fragments (6-9 wks gestation) for 8-11 days, with or without IFN-α/TNF-α. (B) Live-cell microscopy images of co-cultures showing an overlay of phase contrast and GFP immunofluorescence. Tissue fragments with the greatest extent of organoid invasion out of 4-6 fragments are shown. Scale bars: 500 µm. Representative images obtained from other tissue donors are shown in Fig. S4A. (C) Normalized average invasion scores from 4-6 tissue fragments. Images used to score the invasion (scoring table) are shown in Fig. S4B. Boxplot represents median with interquartile range and min-max whiskers (n=5-8 independent experiments, using tissue from different donors). \**P* < 0.05. ns: not significant.

In the untreated EVT condition, we observed clear invasion of GFP-positive trophoblast organoids into the decidual explants (Fig. 5B, Fig. S4A). In contrast, the cytokine-treated EVT organoids, particularly the IFN-α-treated, and undifferentiated organoids showed shallower or no invasion, even though these organoids did attach to the decidual tissue. To draw an unbiased conclusion on the effect of the cytokines on depth of invasion, the images were randomized and manually scored by two independent researchers, blinded for treatment conditions. An ordinal scale between 1 and 4 was used, with a higher value representing deeper invasion, based on a qualitative assessment (Fig. S4B). The normalized average invasion score was significantly reduced for the IFN-α-treated and undifferentiated organoids compared with untreated EVT organoids (Fig. 5C). Moreover, we noticed that the individual tissue fragments co-cultured with IFN-α-treated or undifferentiated organoids never received the highest invasion score of 4 from both researchers, suggesting that deep invasion is impaired in the presence of IFN-α or in the absence of EVT differentiation. For TNF-α, the normalized average invasion score was slightly lower than for the untreated control, but did not reach statistical significance. Altogether, this co-culture system shows that the cytokines, but particularly IFN-α, do not only inhibit EVT invasion into Matrigel, but also in decidual tissue, a situation more closely modeling the in vivo situation.

### Co-culture of trophoblast organoids with first-trimester decidual tissue shows invasion of trophoblasts alongside decidual blood vessels

To better visualize organoid-derived trophoblast invasion into the decidual tissue and investigate potential interactions with blood vessels, the co-cultures were fixed and used for whole-mount immunostaining with antibodies against GFP, endothelial cell marker cluster of differentiation 31 (CD31, encoded by *PECAM1*), and smooth muscle cell marker alpha smooth muscle actin (α-SMA, encoded by *ACTA2*). Many of the tissue fragments contained clear spiral arteries, positive for CD31 and α-SMA (Fig. 6). Organoid-derived, GFP-positive trophoblasts were found invading the decidual stroma, and closely associated with arteries (Fig. 6A-B). Within the stroma, the trophoblasts had rounded morphology (arrowheads in Fig. 6A, GFP-only image), whereas they obtained a more elongated morphology when in contact with blood vessels (arrows in Fig. 6A-B, GFP-only image). For the IFN-α-treated condition, the GFP-positive trophoblasts remained superficial and did not infiltrate the blood vessels or stroma very deeply (Fig. 6C), indicating that the invasion score (Fig. 5C) provides a good indication of EVT invasion in this system.

**Fig. 6.**
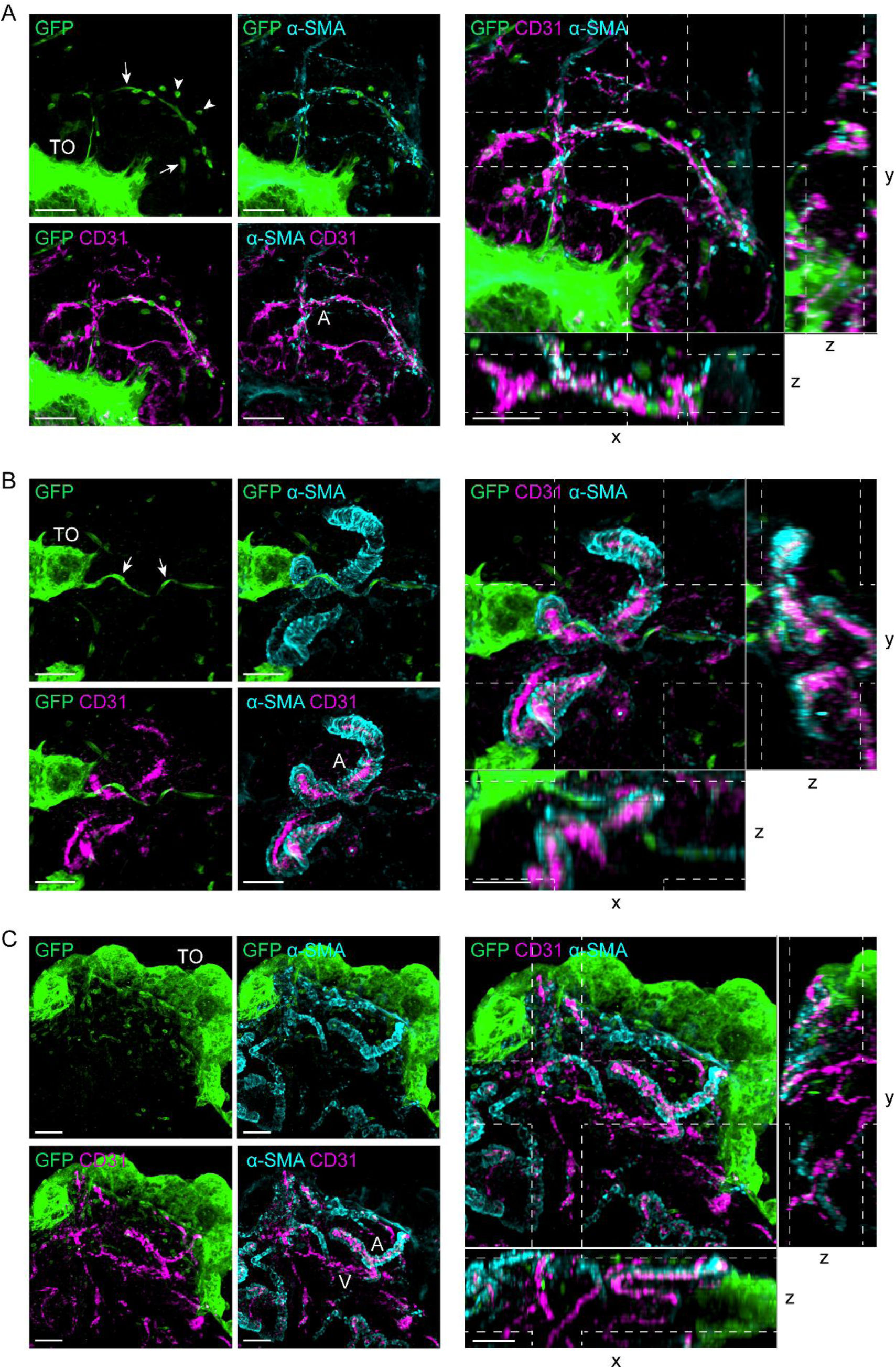
Whole-mount immunostaining of decidua parietalis co-cultured with GFP-positive trophoblast organoids shows invading trophoblasts associating with blood vessels. Maximum intensity projection images showing GFP (GFP reporter-expressing trophoblasts organoids, green) α-SMA (smooth muscle cell marker, cyan), and CD31 (endothelial cell maker, magenta), representative for n=5 independent tissue donors, with the two fragments with highest extent of EVT invasion for each. (A) Untreated organoid condition, showing organoid-derived trophoblasts in decidual stroma and attached to blood vessels. (B) Additional image of the untreated condition, showing close association of organoid-derived trophoblasts with an artery. (C) IFN-α-treated condition, showing shallower trophoblast invasion. Scale bars: 100 µm. x, y, z: dimensions of image stacks. TO: trophoblast organoid; arrows: organoid-derived trophoblasts attached to blood vessels; arrowheads: organoid-derived trophoblasts in decidual stroma (in GFP-only images). A: artery; V: vein (in α-SMA + CD31 overlays).

### Mechanism(s) underlying reduced EVT invasion

Subsequently, we aimed to identify the mechanism(s) underlying the reduced EVT invasion capacity, and investigated whether a molecular pathway targeted by IFN-α and TNF-α could explain the reduced EVT invasion. For this purpose, we FACS-isolated HLA-G-positive EVTs from cytokine-treated and untreated organoids (Fig. S5A), and performed transcriptome sequencing. The viability and percentage of HLA-G-positive cells were not significantly altered by the treatments (Fig. S5B).

For IFN-α-treated EVTs, we identified 325 differentially expressed genes (DEGs) compared with the untreated control (FDR-adjusted *P* < 0.05) (Fig. 7A). For TNF-α-treated EVTs, we identified 22 DEGs compared with the untreated control, of which 16 overlapped with the IFN-α DEGs (Fig. 7A-B). Results of the DEG analysis are provided in Table S1. Gene ontology (GO) analysis indicated an overrepresentation of GO terms related to viral response within the DEGs, as expected from cytokine treatment (Fig. 7C). In line with this, there was a clear enrichment in interferon response gene sets (Fig. 7D). Moreover, there was an enrichment in genes associated with EMT, hypoxia, glycolysis, KRAS and MYC signaling upon IFN-α treatment, which are pathways implicated in invasion (48–52). Among TNF-α-induced DEGs, the glycolysis and MYC targets gene sets were also enriched. These downstream pathways may underly the effect of IFN-α/TNF-α on the invasive capacity of EVTs.

**Fig. 7.**
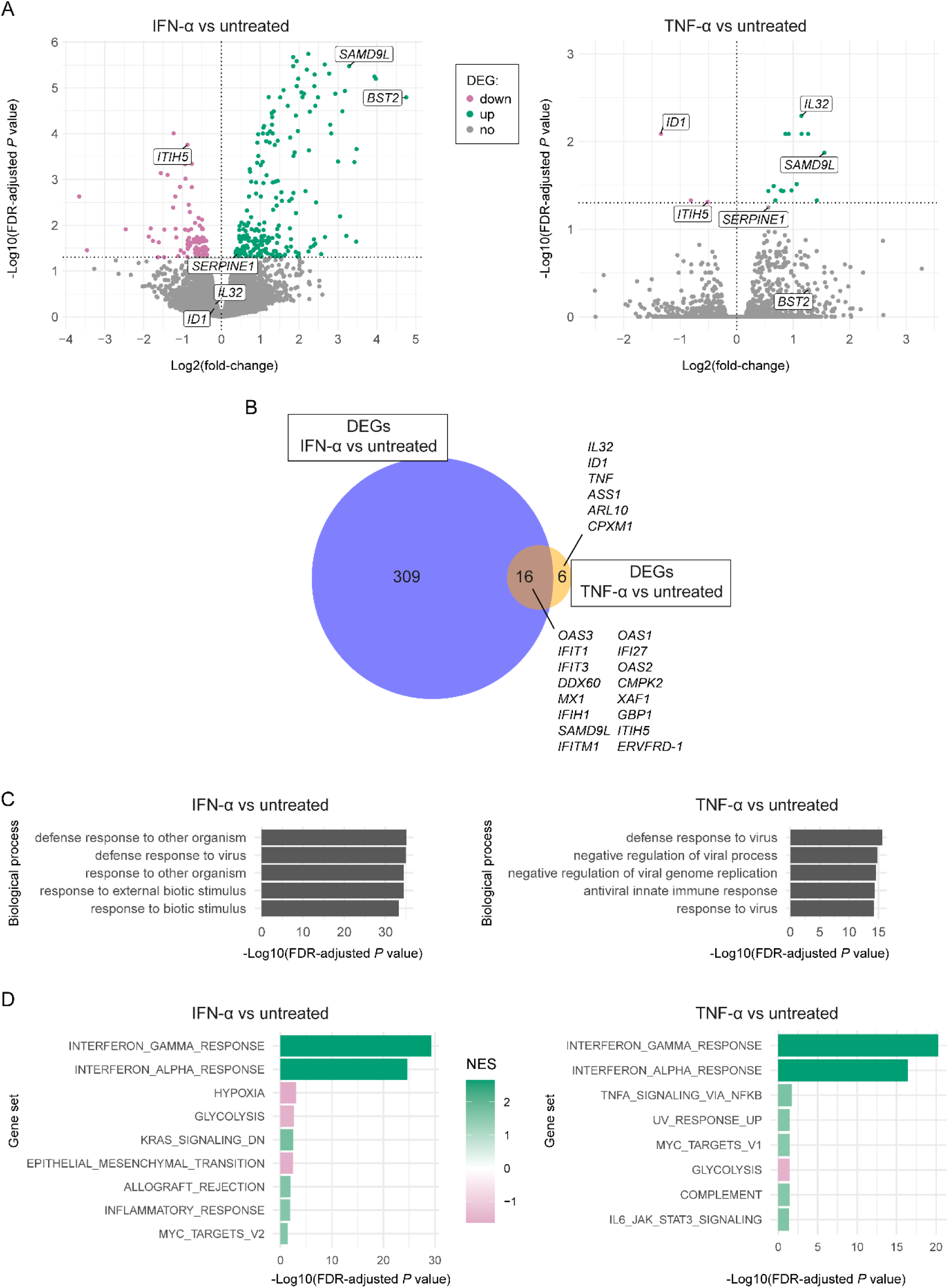
Transcriptome sequencing of EVTs isolated from IFN-α-and TNF-α-treated trophoblast organoids reveals potential mechanisms underlying reduced EVT invasion. Organoids were induced to undergo EVT differentiation for 14 days, with or without IFN-α/TNF-α, and used to isolate HLA-G-positive cells (n=3 independent experiments). (A) Volcano plots showing differentially expressed genes (DEGs) (FDR-adjusted *P* < 0.05) in IFN-α-treated (left) and TNF-α-treated (right) EVTs compared with untreated control EVTs. The candidate genes used for in vitro functional analysis are indicated. (B) Venn diagram showing the overlap between DEGs for IFN-α and TNF-α versus control. Genes specific for TNF-α are sorted by FDR-adjusted *P* value for TNF-α versus control and the list of overlapping genes between IFN-α and TNF-α are sorted by FDR-adjusted *P* value for IFN-α versus control. (C) Top 5 significantly overrepresented biological processes GO terms and (D) significantly enriched gene sets within DEGs. NES: normalized enrichment score.

Because in vitro functional analysis of specific genes may provide further information, we narrowed down the number of candidate genes by overlapping DEGs with the human hallmark EMT gene set (53, 54) as well as matrisome genes, which are associated with the extracellular matrix (55) (Fig. S6A-B). We reasoned that alterations in expression of these genes may influence EVT invasion through an extracellular matrix. In addition, we performed a manual literature search to investigate whether the genes with high fold-change in expression were known to be involved in invasion, and/or associated with pregnancy complications. Based on this, we selected five genes for in vitro functional analysis: *BST2* (IFN-α-specific DEG), *ID1* and *IL32* (TNF-α-specific DEGs), and *ITIH5* and *SAMD9L* (shared DEGs, genes differentially expressed in response to both IFN-α and TNF-α) (Table S2). We also included *SERPINE1*, which was a strong candidate for mediating the TNF-α response (34, 36), although it was not a DEG for TNF-α in our data. Differential expression of the candidate genes is highlighted in Fig. 7A, and their expression is shown in Fig. S6C. Since IL-32 is a cytokine itself, we added it to the organoid medium to test its effect. The other genes were knocked down in TSCs using two different lentivirally expressed shRNAs. Knockdowns of *ID1* and *SAMD9L* resulted in poor cell viability already at the TSC stage, and we could therefore not continue to test their effect on EVT invasion. From the other knockdown TSCs, we generated organoids and induced EVT differentiation in the presence or absence of IFN-α/TNF-α, to assess whether knockdown of the candidate genes mimics or counteracts the IFNA-α/TNF-α-induced phenotype of reduced invasiveness.

As IL-32 was upregulated upon TNF-α treatment, it was hypothesized to mimic the effect of TNF-α. However, addition of IL-32 (10 ng/ml) to the organoids seemed to rather promote EVT invasion (Fig. S7A). For *BST2*, *SERPINE1* and *ITIH5*, we confirmed knockdown on the mRNA level by the two different shRNA sequences (Fig. S7B). As *BST2* was upregulated upon IFN-α treatment, knockdown was hypothesized to rescue IFN-α-reduced EVT invasion. However, due to very strong upregulation upon IFN-α treatment, *BST2* was still upregulated upon IFN-α treatment despite shRNA transduction, and knockdown was therefore not successful under this condition. In the absence of cytokine treatment, *BST2* knockdown slightly reduced the extent of EVT outgrowth, suggesting that this would not counteract the effect of IFN-α (Fig. S7A). *SERPINE1* knockdown was hypothesized to counteract the TNF-α effect and possibly also the IFN-α effect. Independent of cytokine treatment, *SERPINE1* knockdown seemed to increase EVT invasion, although it did not completely prevent the cytokine-induced effect. *ITIH5* was downregulated upon IFN-α and TNF-α treatment, and knockdown was therefore expected to mimic the reduced EVT invasion, independent of cytokine treatment. It indeed decreased the amount of EVT outgrowth, but not to the same extent as IFN-α. Therefore, although these genes had an effect on EVT invasion, they individually did not seem to fully cover the effect of IFN-α and/or TNF-α on EVT invasion.

## Discussion

Immune-mediated inflammatory diseases, in particular SLE, are strongly associated with adverse pregnancy outcomes. This is thought to be caused by disruption of the local immune balance and failure of maternal-fetal tolerance (56). However, the direct effects of the associated pro-inflammatory cytokines on trophoblast development remain poorly understood. In this study, we investigated the effects of IFN-α and TNF-α on EVT differentiation and invasion using multiple model systems. Using different platforms such as TSC monolayer, trophoblast organoids, and first-trimester placental villous explants, we observed that IFN-α and TNF-α inhibit EVT invasion into Matrigel, despite normal EVT differentiation. In a co-culture of trophoblast organoids with first-trimester decidual explants, particularly IFN-α inhibited EVT invasion into decidua in the same way as in Matrigel. Transcriptome sequencing on the EVT subpopulation from organoids suggested that IFN-α and TNF-α affected several pathways implicated in invasion.

EVT differentiation and invasion into the maternal decidua are essential placentation processes, taking place predominantly during the first trimester (3, 7). Within the decidua, EVTs remodel spiral arteries into large-diameter vessels with reduced contractility, which enables proper maternal blood flow to the intervillous space. Incomplete EVT invasion and spiral artery remodeling can lead to discontinuous and high-rate blood flow, causing oxidative stress, damaging of the placental villous tree, and disturbed maternal-fetal exchange, as seen in early-onset preeclampsia (57). As our results indicate that IFN-α and TNF-α directly inhibit EVT invasion, this mechanism may contribute to the increased incidence of preeclampsia and other adverse outcomes in SLE pregnancies.

To be able to thoroughly study EVT invasion in a physiological way, we used two assays: a single-organoid system, and a co-culture system of GFP-expressing trophoblast organoids with first-trimester decidual explants. The single-organoid assay allowed quantification of EVT invasion into extracellular matrix, while the co-culture assay enabled qualitative assessment of EVT interaction with decidual tissue. GFP reporter-expressing TSCs allowed visualization of EVTs within different decidual compartments, such as stroma, blood vessels and glands, using whole-mount immunostaining. The presence of these structures and the three-dimensionality make this a more physiological and translational model of EVT invasion than for example the widely-used transwell invasion assays, and other in vitro culture systems out of their tissue context. Moreover, since TSCs can be genetically manipulated, both assays offer excellent opportunities to study the role of specific genes in EVT invasion and early placenta formation.

Within the decidual tissue, we observed organoid-derived trophoblasts in the stroma, and in close contact with blood vessels. This suggests that EVTs are attracted by various decidual cell types in this system, similarly as in vivo (58). This observation also highlights the potential of this system for studying spiral artery remodeling, although the currently maximal time window of the co-culture might be a limiting factor. Spiral artery remodeling takes place between 6 and 20 weeks of gestation, with individual arteries undergoing remodeling asynchronously, but the duration for individual arteries is unknown (6). This would be worth further investigation with further optimization of the co-culture system and high resolution imaging. Additional characterization would also allow studying EVT association with, for example, decidual immune cells, glandular epithelium and lymphatics. Moreover, spatial omics techniques could be used to determine whether organoid-derived EVT localization correlates with altered gene expression, and how this compares to in vivo profiles of EVT subtypes as determined by others (59). Limitations of the co-culture system that we experienced include dependency on donor tissue availability, occasional poor tissue quality, and technical complexity, although these are all challenges that can be overcome. In addition, high inter-donor and inter-experimental variability in terms of invasion depth required multiple repetitions of the experiment and normalization. Nevertheless, the co-culture system opens up many possibilities for future research.

An important question is whether the cytokine concentrations used here reflect those in patients. In general, cytokines primarily function via paracrine and autocrine signaling to regulate local immune responses, and systemic levels are therefore low (60). In SLE and other immune-mediated inflammatory diseases, however, systemic levels are elevated, especially during disease flares (61, 62). Local cytokine concentrations are probably more relevant for EVT invasion, but they are difficult to measure. For the choice of concentrations, we relied on concentrations used in the literature. In the co-culture system, decidual cells are also exposed to the added cytokines, and most likely consume them (63), possibly interfering with results. This may explain the weaker effect of TNF-α in co-culture with decidua than in culture of organoids alone. Another explanation would be that the co-culture system may not be sensitive enough to detect subtle differences, as many factors are involved.

To identify target genes and pathways by which IFN-α/TNF-α reduce EVT invasion, we performed transcriptome sequencing of HLA-G-positive EVTs, isolated from cytokine-treated organoids. Pathway analysis indicated an enrichment for genes related to EMT, hypoxia, glycolysis, KRAS and MYC signaling, which are all involved in invasion (48–52). IFN-α-/TNF-α-induced dysregulations of these pathways may therefore underlie the observed phenotype. During the preparation of this manuscript, Simoni *et al*. reported that IFN-β, another type I interferon, inhibits EVT invasion in an organ-on-a-chip model, using primary EVTs and endothelial cells (28). Consistent with our results, they showed that addition of IFN-β decreases EVT invasion through Matrigel towards endometrial endothelial cells, which is accompanied by alterations in EMT-related gene expression. However, that study did not distinguish between direct and indirect effects on EVTs, as both endothelial cells and EVTs were exposed to IFN-β. In addition, the absence of other decidual cells, such as stromal and immune cells, did not allow for studying the interplay with other cell types. Our study using organoid and co-culture models is therefore a valuable follow-up, and the results being in line provides further evidence for the role of type I interferons in EVT invasion.

Although our transcriptome data did not reveal clear target genes to explain the reduced invasion, we selected some DEGs for functional analysis, based on their involvement in EMT or regulation of the extracellular matrix, as well as known association with invasion in the literature. IL-32 had an opposite effect of TNF-α and promoted EVT invasion, which is in line with findings of a study using HTR-8/SVneo cells (64). The TNF-α-induced upregulation of *IL32* may therefore reflect a compensatory mechanism. Since IL-32 also upregulates *TNFA* (65), these cytokines might form a negative feedback loop regulating EVT invasion. Knockdown of the other candidate genes, *BST2*, *SERPINE1* and *ITIH5,* altered the extent of EVT outgrowth, but the individual genes could not fully explain the reduced invasiveness. Since there were many other DEGs in our data, we might simply not have selected the most important ones. However, the phenotype could also be a combined effect of multiple genes, or we might have missed key targets due to the timing of sampling, indirect effects via cytotrophoblasts or syncytiotrophoblasts, or involvement of non-transcriptional mechanisms. Further research is therefore needed to identify direct mediators.

In pregnancy, type I IFNs and TNF-α at the maternal-fetal interface are known to play a role in protecting the fetus against bacterial and viral infection (26). Considering their effect on EVT invasion, secretion of low levels of these cytokines by decidual cells might play an additional role in preventing excessive invasion into the uterine wall. Together with other signals, they may balance invasion depth. Interestingly, when interstitial EVTs undergo terminal differentiation into placental bed giant cells in vivo, they upregulate type I IFN receptors (59). As the formation of giant cells is thought to be a mechanism preventing excessive invasion (66), it is conceivable that type I IFN signaling plays a role in that. It would be interesting to further investigate the involvement of these cytokines in conditions such as placenta accreta, where the placental villi and EVTs invade the myometrium (67). Beyond immune-mediated inflammatory diseases, the effect of these cytokines on EVT invasion may also contribute to the increased risk for adverse pregnancy outcomes upon infection and in other chronic inflammatory conditions such as obesity (68–70).

Women with active or recently-diagnosed SLE are advised to postpone pregnancy until disease activity is low or in remission (71). A deeper understanding of the mechanisms driving pregnancy complications in SLE and other immune-mediated inflammatory diseases may provide a rationale for such management as well as for medical treatment decisions. Moreover, it may reveal new targets for preventative treatments. This work improves our understanding of early placenta development, and offers mechanistic insights into the increased risk for adverse pregnancy outcomes in SLE patients.

## Methods

### Human tissue collection

First-trimester placental and decidual tissue were obtained from the Human Immune System Mouse Facility of the Amsterdam University Medical Center, Amsterdam, and de-identified prior to use in the current study. The collection of extra-embryonic tissue was approved by the Amsterdam University Medical Center Biobank Review Committee (AMC 2016_246). Alternatively, decidual tissue was collected from the Leiden University Medical Center, as approved by the Medical Ethical Committee of the Leiden University Medical Center (B21.054). The human tissue was donated for scientific research with written informed consent from donors undergoing elective abortion without medical indication.

### TSC and organoid culture and differentiation

Culture and differentiation of TSCs and trophoblast organoids were performed as described in our previous work (46). In short, human TSCs, isolated from first-trimester placental tissue, were cultured and induced to undergo STB and EVT differentiation in monolayer following protocols of Okae *et al*. (41). Trophoblast organoids were generated by seeding TSCs into domes of Matrigel growth-factor reduced basement membrane matrix (Corning, 356231), and organoid culture and EVT differentiation were performed according to protocols established by others (42, 43).

### Placental villous explant culture

First-trimester placental tissue was washed in PBS. Placental villi were dissected and seeded into the center of wells of a 48-wells plate pre-coated with 50 µl of Matrigel, after which they were spread using a needle (5 fragments per condition). To allow attachment to the Matrigel, the explants were overlaid with 20 µl DMEM/F-12 + GlutaMAX (Gibco, 31331-028) with 1% penicillin-streptomycin, and incubated at 37 °C in 5% CO_2_ and 5% O_2_ for 4 h or overnight. Subsequently, 250 µl medium including treatment was added, and the explants were incubated for 5-7 days, with medium refreshments every 2-3 days.

### Cytokine treatment

Human recombinant TNF-α (PreproTech, AF-300-01A) or IFN-α 2A (STEMCELL Technologies, 78076) were added to the medium in a concentration of 10 ng/ml and 5 ng/ml, respectively. Human recombinant IL-32β (R&D Systems, 6769-IL-025) was added in a concentration of 10 ng/ml. These concentrations were based on concentrations used in the literature, and were not toxic to our TSC lines.

### Lentiviral transduction

To generate green fluorescent protein(GFP)-expressing TSCs, 5×10^4^ cells were transduced with lentivirus 24 h after seeding. The lentivirus was produced in-house using standard packaging protocols with the MISSION® pLKO.1-puro-CMV-TurboGFP Positive Control Plasmid (Merck, SHC003). Forty-eight hours post-transduction, cells with high GFP fluorescence were isolated using a Sony SH800 cell sorter, and maintained as TSC monolayer culture. To generate knockdowns, TSC were transduced with short hairpin RNA (shRNA) vectors as described previously (46), using the shRNA sequences presented in Table S3.

### RNA isolation and quantitative real-time PCR (RT-qPCR)

RNA isolation, cDNA synthesis and RT-qPCR of TSCs and organoids were performed as described in our previous work (46). To release organoids from the Matrigel, the domes were incubated with 2 mg/ml dispase II (Sigma-Aldrich, D4693), for 2 x 5 min at 37 °C, with resuspending in between incubations using a 1% BSA-coated tip. The organoids were then washed in 1 mM EDTA/PBS, and subsequently used for lysis. Primer sequences used for RT-qPCR are provided in Table S4.

### Immunofluorescence staining and microscopy of organoids

For immunofluorescence imaging, organoid domes were fixed in 2% PFA/PBS for 30 min at 4 °C, and immunostained within the remaining Matrigel. Permeabilization and blocking was done in 0.5% Triton X-100 + 4% BSA in PBS for at least 4 hours or overnight at 4 °C. Primary antibodies as well as secondary antibodies with 2 μg/ml DAPI were diluted in Bright Diluent (Immunologic, VWRKUD09), and both stainings were executed for 2 nights at 4 °C. See Table S5 for antibody dilutions and product information. Washings were done in 0.1% Triton X-100 + 0.2% BSA in PBS for 3 x 30 min at room temperature. Organoids were cleared in 50% glycerol/dH2O + 2.5 M fructose, and imaged on a Leica Thunder Imager DMi8 inverted wide-field microscope. To enhance contrast, grayscale values were adjusted in Leica Application Suite software. All settings and adjustments were kept equal between conditions.

### Single-organoid EVT invasion assay

To quantify EVT invasion from individual organoids, organoids were generated within Matrigel domes and cultured for 5 days in trophoblast organoid medium (TOM) and 5 days in trophoblast organoid(TO)-EVT medium A (see previous work for medium compositions (46)). Subsequently, organoids were released from the Matrigel domes by incubation in Cell Recovery Solution (Corning, 354253) for 1 h on ice. Using a stereo microscope and 0.1% BSA-coated pipette tips, single organoids were then transferred into the center of wells of a 96-wells plate pre-coated with 40 µl Matrigel (10 organoids per condition per independent experiment). The organoids were overlaid with TO-EVT medium A, in which they were cultured for 2 extra days, and then switched to TO-EVT medium B for another 7 days. Incubation was done at 37 °C in 5% CO_2_, and medium was refreshed every 2-3 days. Organoids were stained with 4 µM Calcein AM (eBioscience, 650853) for 30 min at 37°C. Z-stack images (10 µm steps) were taken of the single organoids, including all invading EVTs, using a Thunder Imager DMi8 wide-field fluorescence microscope (Leica), and analyzed using Imaris software (Oxford Instruments-Andor Technology, v10.1.1).

### Trophoblast organoid-decidua explant co-culture

Decidua parietalis tissue (6-9 wks gestation) was washed in PBS and dissected into ∼1-3 mm cubes under a stereomicroscope. These fragments were either immediately used, or stored in Cell Banker I (AMSBIO, 11910) in liquid nitrogen and thawed on the starting day of co-culture.

GFP-transduced TSCs were used to generate organoids in Matrigel domes for 5 days in TOM and induced to undergo EVT differentiation for 7-9 days. Organoids were then released from the Matrigel using Cell Recovery Solution for 20 min on ice, and resuspended in TO-EVT medium B with 50 µg/ml gentamicin (Gibco, 15710-049). The decidual tissue fragments were placed onto a layer of 200 µl Matrigel in wells of a 4-wells plate (Thermo Scientific, 176740) (4-6 fragments per well), if possible with the luminal epithelial side facing upwards, and evenly distributed using a needle. The plate was incubated for 25 min at 37 °C in 5% CO_2_ and 5% O_2_ to allow the Matrigel to set, followed by another 20 min upside-down. Next, the organoid suspension was added on top of the Matrigel layer. Alternatively, DMEM/F12 + GlutaMAX with 1% pen/strep and 50 µg/ml gentamicin was first added, and replaced by the organoid suspension in TO-EVT medium B later that day. The medium was refreshed every 2-3 days and the co-culture was analyzed after 8-11 days.

### Immunofluorescence staining and microscopy of decidual tissue

Co-culture were washed twice with PBS, after which decidual fragments were transferred to individual wells of a 48-wells plate using tweezers. They were fixed and permeabilized in 4% PFA/PBS with 1% Triton X-100 for 1 h at room temperature, after which they were washed 3 x 20 min in PBS at room temperature and stored at 4 °C until further processing.

For whole-mount immunostaining in a 48-wells plate, non-specific binding sites of the tissue fragments were first blocked by incubation in 1% BSA/PBS with 1% Triton X-100 and 0.1% saponin (Acros Organics, 419231000) (blocking/permeabilization buffer) overnight at 4 °C. The next day, primary antibodies, diluted in blocking/permeabilization buffer, were added to the tissue fragments to incubate 3 nights at 4 °C on a shaking platform. See Table S5 for antibody dilutions. Secondary antibody incubation (optionally in the presence of 2 μg/ml DAPI) was done for 2 nights under the same conditions. Washing was done in between and after incubation steps in PBS with 0.2% Triton -X-100 for 6 x 30 min at room temperature. Subsequently, tissue fragments were transferred to 1.5 ml Eppendorf tubes and incubated 2 x 30 min in 100% ethanol. Next, they were cleared in ethyl cinnamate (Sigma-Aldrich, W243000) for 2 days on a shaking platform, and kept at room temperature.

Using a micro spatula, the cleared samples were transferred to a 96-wells black/clear bottom plate (Thermo Scientific, 10281092) with 150 µl ethyl cinnamate per well. Whole samples were imaged on an Andor Dragonfly 500 spinning disk confocal microscope with 3-6 µm step size. Images were adjusted using ImarisViewer software (Oxford Instruments-Andor Technology, v10.2.0).

### Isolation of EVTs from organoids

To be able to perform RNA sequencing on the EVT fraction of the EVT-differentiated organoids, we isolated the HLA-G-positive cells. Organoids were induced to undergo EVT differentiation for 14 days, after which they were released from the Matrigel domes using dispase II as described above. To make the organoids single cell, they were incubated in TrypLE (Gibco, 12604) for 5 min at 37 °C, and filtered through a 70 µm filter. The flow-through cells were stained with FITC-conjugated anti-HLA-G antibody (EXBIO, 1F-292-C100, 1:200) or FITC-conjugated mouse IgG1 isotype control (BD, 555748, 1:200) in PBS supplemented with 0.05% BSA and 0.01% NaN_3_ for 15 min at room temperature, and with Bioscience Fixable Viability Dye eFluor 780 (Invitrogen, 65-0865-14) in PBS for another 15 min. The HLA-G-positive cell fraction was identified based on the isotype control and collected using a FACSAria Ilu SORP flow cytometric cell sorter (Becton Dickinson). Doublets and dead cells were excluded, and data were analyzed using FlowJo software (v10). The isolated cells were lysed and used for RNA isolation.

### Transcriptome sequencing

Total RNA libraries were prepared using the KAPA mRNA HyperPrep kit (Roche, 08098123702), and 150 bp paired-end sequencing was performed on an Illumina NovaSeq X Plus system. Quality control, mapping and data analysis were performed as previously described (46), with the following alterations in version numbers: Ensembl (v107), RSeQC (v5.0.1), MultiQC (v1.13), R programming language (v4.4.1), EdgeR Bioconductor package (v4.2.2), sva package (3.54.0). Gene set enrichment analysis was performed using the fgsea Bioconductor package (v1.32.4), and Venn diagrams were created using the VennDiagram package (1.7.3).

### Statistics

Statistical analyses were performed in GraphPad Prism (v10.2.0) or R programming language v(v4.4.1). For RT-qPCR data, a one-way ANOVA with Dunnett’s multiple comparisons test was performed, and an additional Bonferroni correction was done to adjust for testing multiple genes. Invasion quantification data were analyzed using a generalized linear model with Poisson distribution for count data, and Gaussian distribution for distance measurements. For semi-quantitative invasion score data, a Kruskall-Wallis test with Dunn’s multiple comparisons test was performed. *P* < 0.05 was considered statistically significant.

### Data availability

Transcriptome sequencing data have been deposited in the Gene Expression Omnibus database under accession number GSE302798.

## Funding

This work was supported by the Novo Nordisk Foundation (NNF21CC0073729, reNEW to SMCSL).

## Acknowledgments

We would like to thank the donors for the placental and decidual tissue that made this study possible. We would also like to thank present and past lab members of the Chuva de Sousa Lopes and Afink groups for fruitful discussions and technical advice. This work made use of the Dutch national e-infrastructure with support of the SURF Cooperative using grant no. EINF-14364.

## Notes

https://www.ncbi.nlm.nih.gov/geo/query/acc.cgi?acc=GSE302798

### Competing Interest Statement

The authors have declared no competing interest.

