## Supplementary Material for "The pro-inflammatory cytokines IFN-α and TNF-α inhibit organoid-derived extravillous trophoblast invasion"

### Supplementary Figures

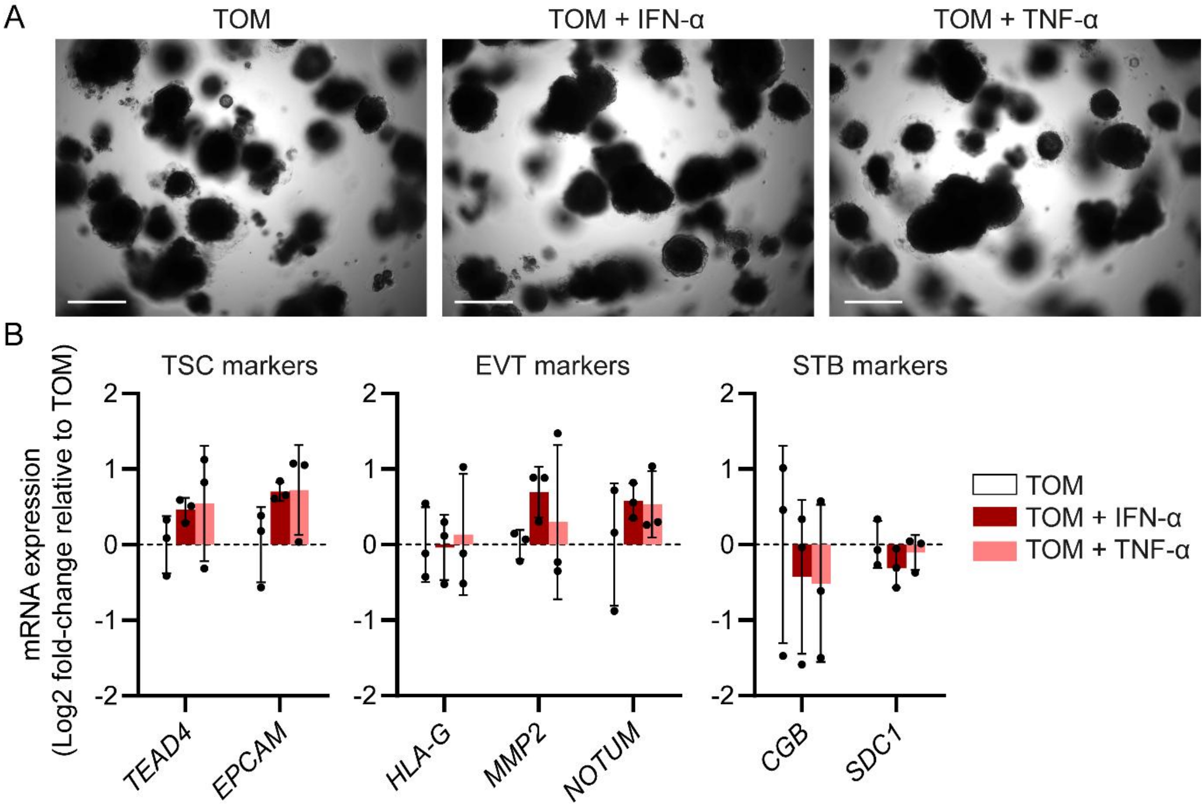


**Fig. S1. IFN-α and TNF-α do not affect trophoblast organoid formation.** Organoids were grown in Matrigel domes in trophoblast organoid medium (TOM) for 5 days, followed by another 14 days in the presence or absence of IFN-α (5 ng/µl) or TNF-α (10 ng/µl). (A) Representative phase-contrast images. Scale bars: 400 µm. (B) Expression of TSC, EVT and STB markers on the mRNA level, measured by RT-qPCR. Bars represent mean Log2 fold-change ± SD relative to the mean of untreated organoids (n=3 independent experiments). All results were non-significant compared with the untreated TOM condition.

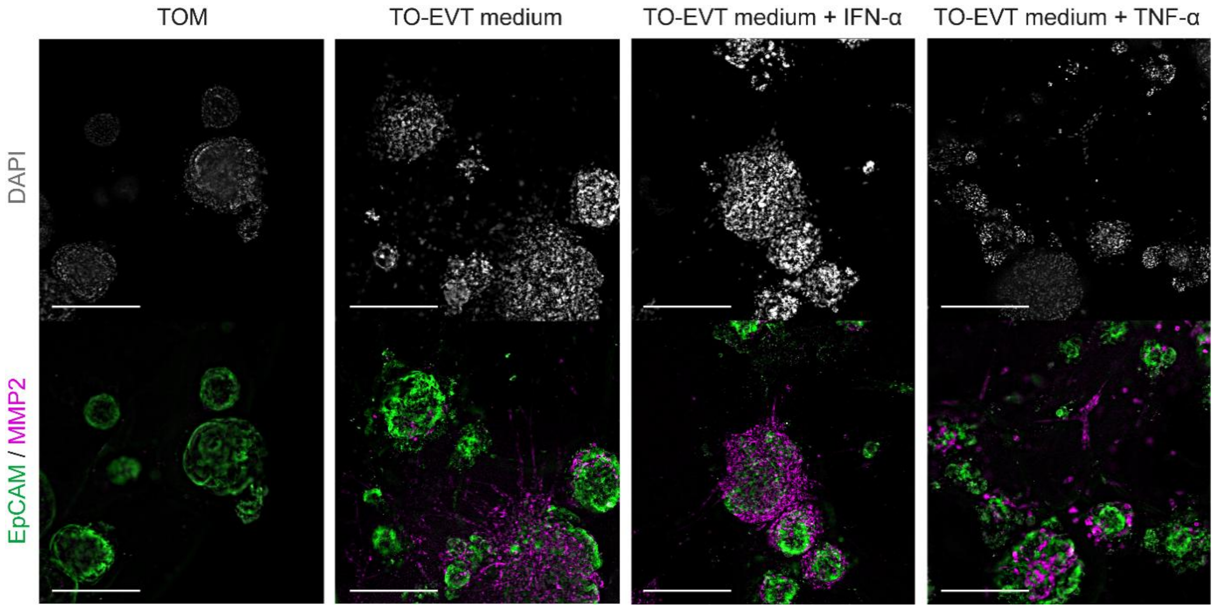


**Fig. S2 (related to Fig. 2). Immunofluorescence images showing that IFN-α and TNF-α inhibit EVT invasion but do not affect EVT differentiation in trophoblast organoids.** Organoids at EVT differentiation day 14 were stained for EpCAM (TSC marker, green), MMP2 (EVT marker, magenta), and DAPI (nuclei, white). Scale bars: 400 µm.

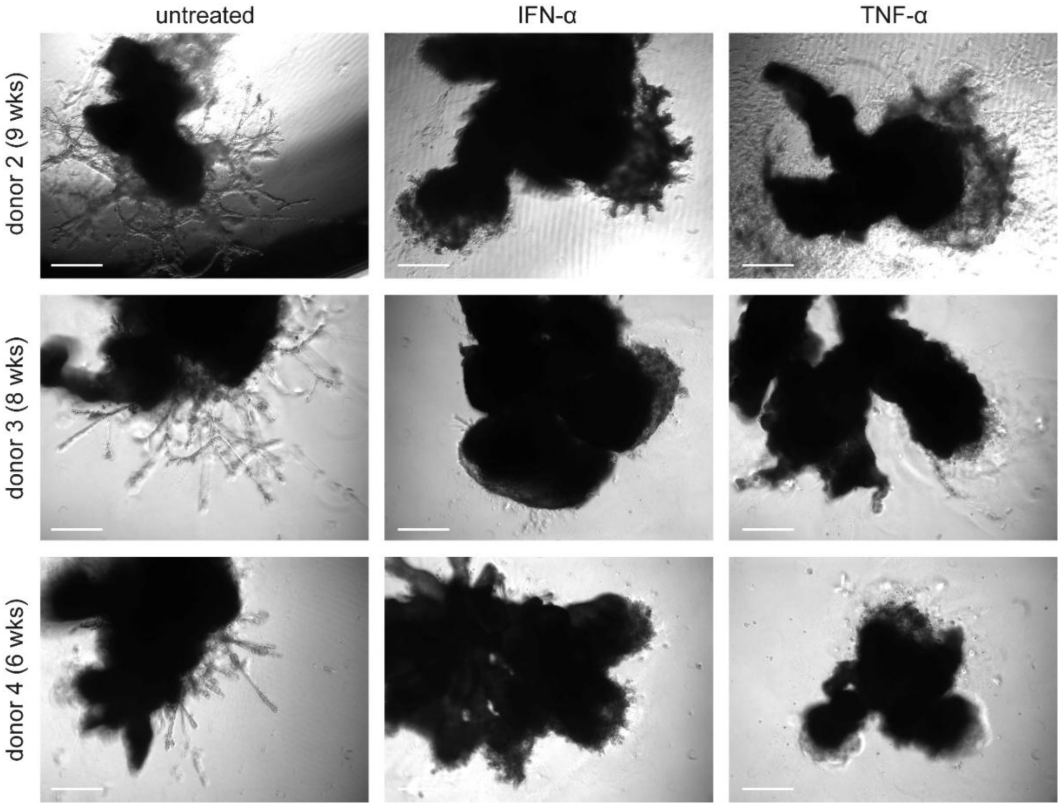


**Fig. S3 (related to Fig. 3). Experiments with additional tissue donors showing that IFN-α and TNF-α affect EVT invasion in placental villous explants.** Scale bars: 400 µm.

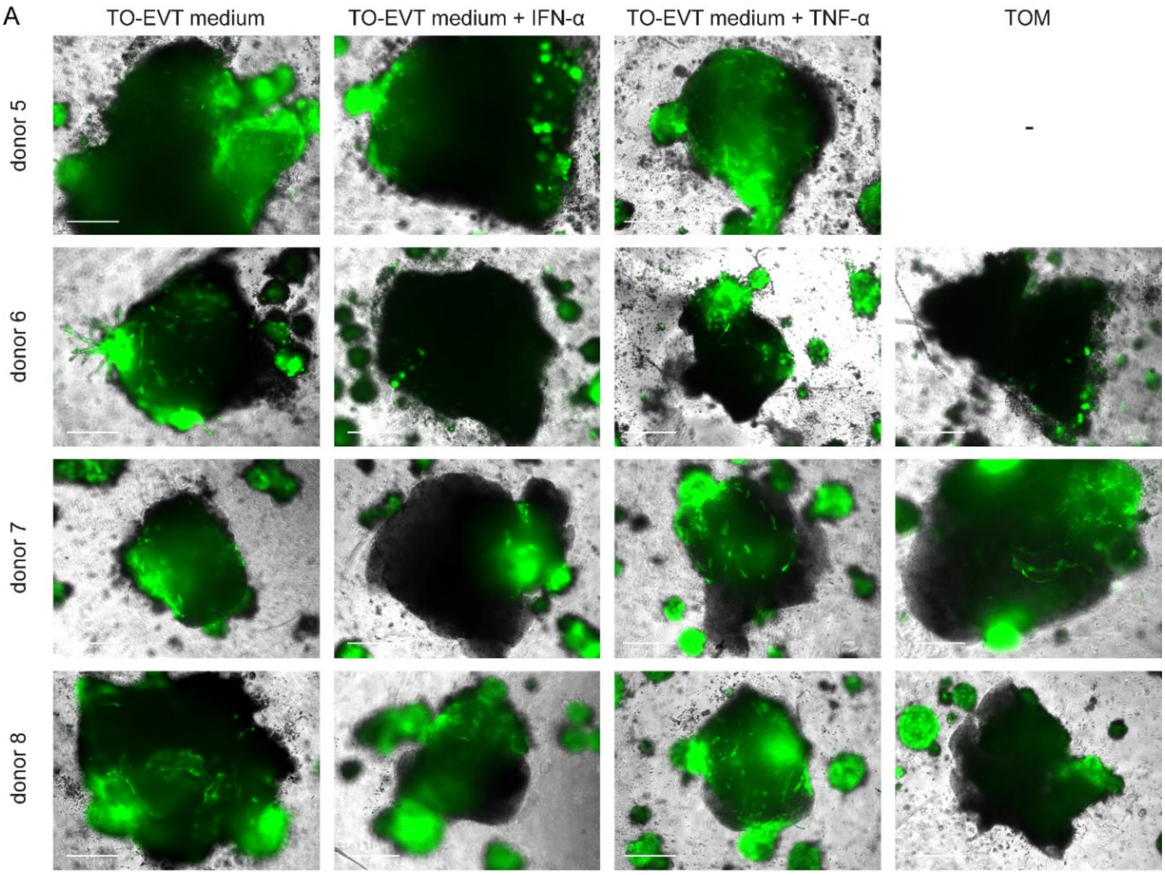


B

| Invasion score | | Example images |
|---|---|---|
| 4 | Very clear/deep invasion | 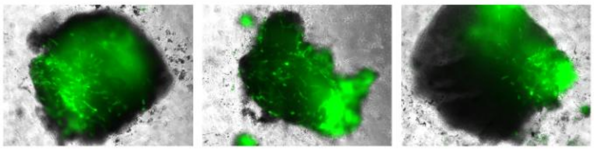 |
| 3 | Clear invasion | 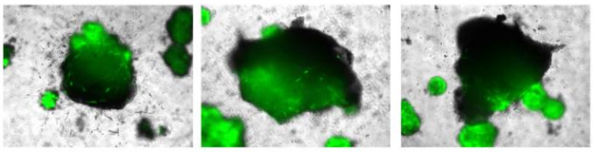 |
| 2 | (Possibly) some invasion | 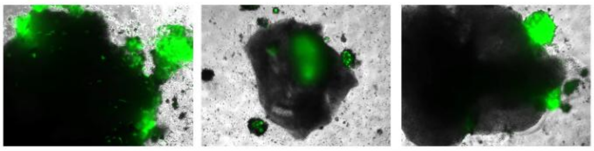 |
| 1 | (Probably) no invasion | 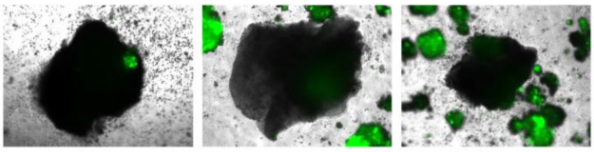 |

**Fig. S4 (related to Fig. 5). Co-culture of GFP-positive trophoblast organoids and first-trimester decidua parietalis.** (A) Experiments with additional tissue donors. Scale bars: 500 µm. (B) Invasion scoring table used to semi-quantify organoid-derived EVT invasion into decidual explants.

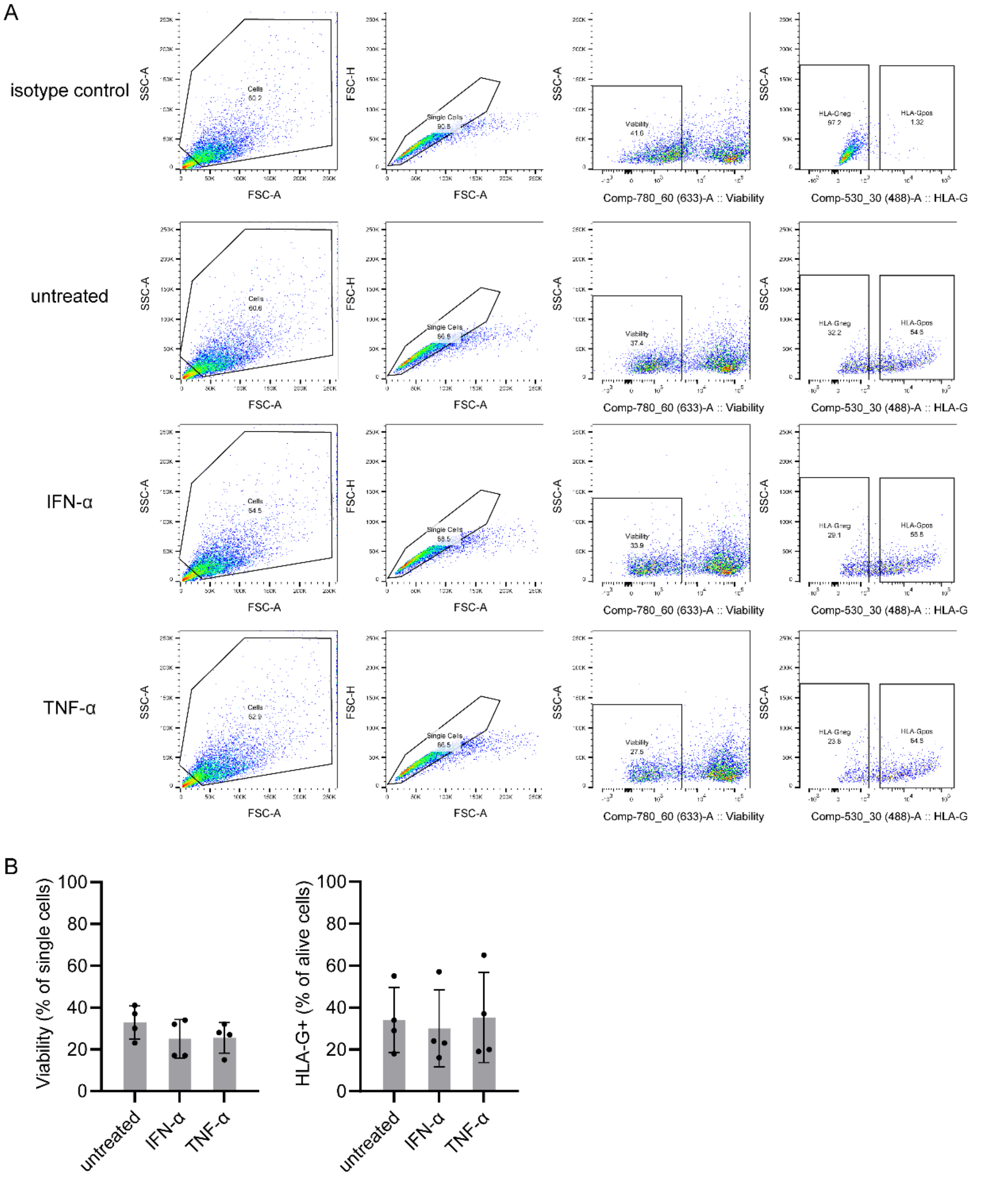


**Fig. S5 (related to Fig. 7). Isolation of HLA-G-positive cells by FACS.** (A) Gating strategy used. The images are representative for n=4 independent experiments. For the isotype control condition, a pool of untreated, IFN-α- and TNF-α-treated cells was used. (B) Percentages of viable and HLA-G-positive cells. Bars represent mean ± SD of n=4 independent experiments.

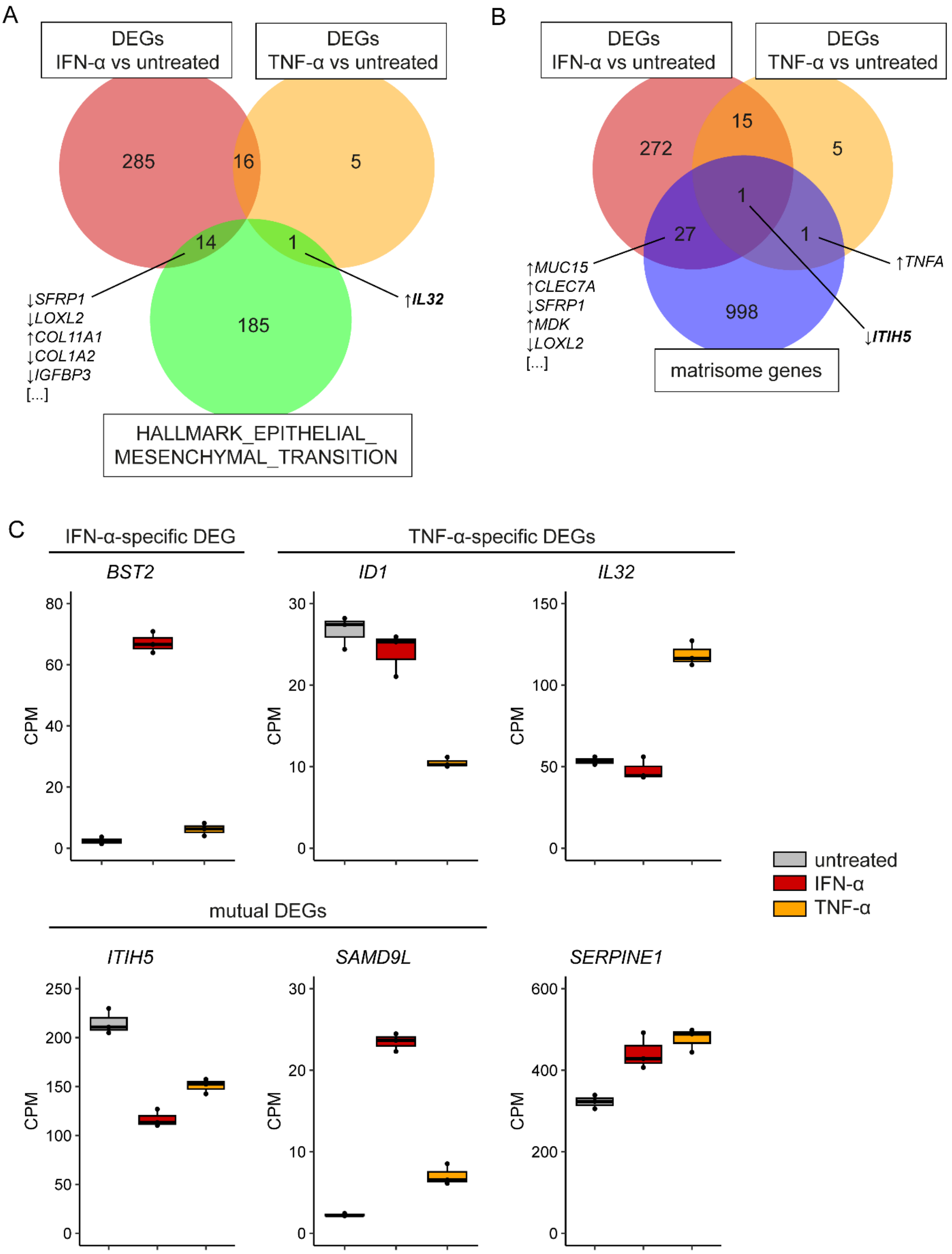


**Fig. S6 (related to Fig. 7). Selection of candidate genes for in vitro functional analysis.** Venn diagrams (using annotated gene symbols) overlapping the IFN-α- and TNF-α-induced DEGs with (A) EMT genes (1, 2) and (B) matrisome genes (3). The indicated genes are sorted by FDR-adjusted *P* value, and arrows indicate up- or downregulation compared with the untreated control. (C) Expression of candidate genes in transcriptome data of HLA-G+ EVTs isolated from organoids, as batch-corrected counts per million (CPM). Boxplots represent median with interquartile range and min-max whiskers (n=3 independent experiments).

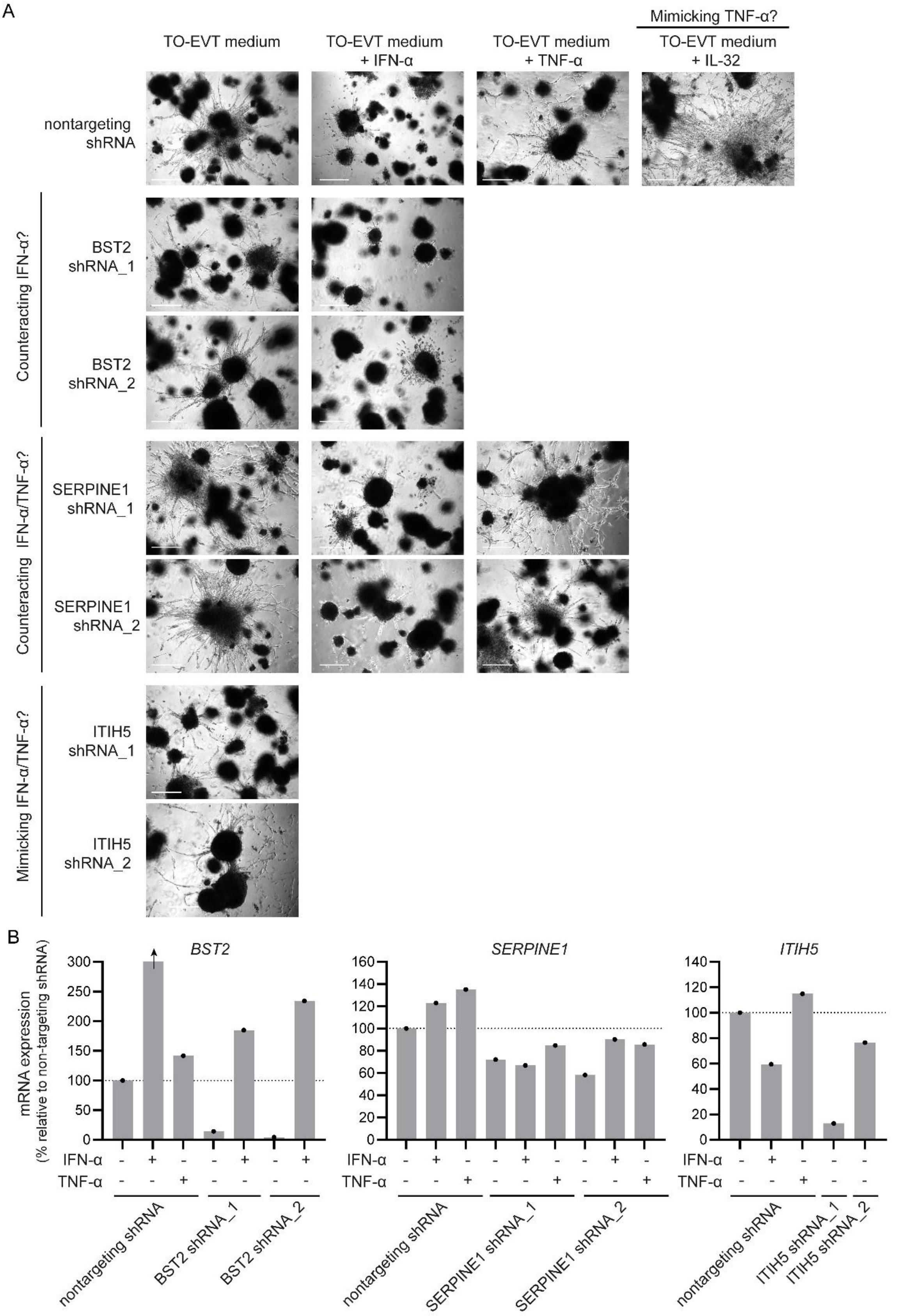


**Fig. S7. Candidate genes mimicking/rescuing EVT invasion in organoids.** TSCs transduced with shRNA were used to generate organoids, which were induced to undergo EVT differentiation for 14 days in the presence or absence of cytokines. In addition, organoids were treated with IL-32

(10 ng/ml). (A) Representative phase-contrast images (n=1 for knockdowns, and n=3 independent experiments for IL-32). Scale bars: 500 µm. (B) Candidate gene mRNA expression as percentage relative to the untreated nontargeting shRNA condition, measured by RT-qPCR. For the nontargeting shRNA + IFN-α condition, *BST2* expression was 1278% relative to the control, exceeding the y-axis (indicated by arrow).

### Supplementary Tables

**Table S1. Results of DGE analysis from EVT fraction of IFN-α/TNF-α-treated organoids compared with untreated controls.** See separate Excel file.

**Table S2. Selection of candidate genes (related to Fig. 7 and Fig. S6-7).**

| Gene symbol | Gene name | Log2(FC) relative to untreated control | | EMT gene set (1, 2) | Matrisome (3) | Literature invasion | Other |
|---|---|---|---|---|---|---|---|
| | | IFN-α | TNF-α | | | | |
| *BST2* | bone marrow stromal cell antigen 2 | 4.76* | 1.27 | no | no | trophoblast (4, 5) | link with PE (5) |
| *ID1* | inhibitor of DNA binding 1 | -0.18 | -1.34* | no | no | trophoblast (6) | |
| *ITIH5* | inter-alpha-trypsin inhibitor heavy chain 5 | -0.87* | -0.51* | ECM regulator | no | other cell types (7, 8) | |
| *IL32* | interleukin 32 | -0.16 | 1.15* | no | yes | trophoblast (9) | link with PE (9, 10) |
| *SAMD9L* | sterile alpha motif domain containing 9 like | 3.29* | 1.56* | no | no | other cell types (11, 12) | |
| *SERPINE1* | serpin family E member 1 (also known as PAI-1) | 0.42* | 0.56 | ECM regulator | yes | trophoblast (13, 14) | |

Green shading: reason to select. FC: fold-change. EMT: epithelial–mesenchymal transition. *FDR-adjusted $P < 0.05$ compared with untreated control. ECM: extracellular matrix. PE: preeclampsia. (1-14): references.

**Table S3 (related to Methods). shRNA sequences used for knockdown of candidate genes.**

| shRNA | Manufacturer | Cat. no. | Sequence |
|---|---|---|---|
| BST2 shRNA_1 | Merck | TRCN0000107016 | CCAGGTCTTAAGCGTGAGAAT |
| BST2 shRNA_2 | Merck | TRCN0000107019 | GAGGGAGAGATCACTACATTA |
| ITIH5 shRNA_1 | Merck | TRCN0000118263 | CCGTTTCAGTATCATTGGATT |
| ITIH5 shRNA_2 | Merck | TRCN0000118265 | CTGCATCTTCACCATTGGCAT |
| SERPINE1 shRNA_1 | Merck | TRCN0000052270 | GCATCTGTACAAGGAGCTCAT |
| SERPINE1 shRNA_2 | Merck | TRCN0000052271 | CAGACAGTTTCAGGCTGACTT |
| nontargeting shRNA | Merck | SHC002 | CCGGCAACAAGATGAAGAGCACCAACTCG AGTTGGTGCTCTTCATCTTGTTGTTTTT |

**Table S4 (related to Methods). Primer sets used for quantitative real-time PCR.** *EEF2* and *GUSB* were used as reference genes.

| Target gene symbol | Forward primer | Reverse primer | Amplicon size (bp) |
|---|---|---|---|
| *BST2* | TCTCCTGCAACAAGAGCTGACC | TCTCTGCATCCAGGGAAGCCAT | 113 |
| *CGB* | CAGCATCCTATCACCTCCTGGT | CTGGAACATCTCCATCCTTGGT | 102 |
| *EEF2* | CTGGAGATCTGCCTGAAGGA | CGACCGGGTCAGATTTCTT | 70 |
| *EPCAM* | GCCAGTGTACTTCAGTTGGTGC | CCCTTCAGGTTTTGCTCTTCTCC | 122 |
| *GUSB* | GGAGTGCAAGGAGCTGGAC | ATTGAAGCTGGAGGGAACTG | 145 |
| *HLA-G* | CCACCACCCTGTCTTTGACTAT | ACGTCCTGGGTCTGGTCCT | 114 |
| *ITGA1* | CCGAAGAGGTACTTGTTGCAGC | GGCTTCCGTGAATGCCTCCTTT | 107 |
| *ITIH5* | CAGTCACTCCAGACAGCATCAG | CCACGTACTTGTTGAGGAGCCT | 120 |
| *MMP2* | TGGCACCCATTTACACCTACAC | ATGTCAGGAGAGGCCCCATAGA | 91 |
| *NOTUM* | CTACTGGTGGAACGCAAACATGG | CGCACCACCTCCTGGATGATG | 129 |
| *SDC1* | CTATTCCCACGTCTCCAGAACC | GGACTACAGCCTCTCCCTCCTT | 102 |
| *SERPINE1* | CTCATCAGCCACTGGAAAGGCA | GACTCGTGAAGTCAGCCTGAAAC | 154 |
| *TEAD4* | CAGGTGGTGGAGAAAGTTGAGA | GTGCTTGAGCTTGTGGATGAAG | 120 |

**Table S5 (related to Methods). Primary and secondary antibodies used for immunofluorescence.**

| Antibody | Manufacturer | Cat. no. | Source | Clone | Dilution |
|---|---|---|---|---|---|
| anti-goat IgG Alexa Fluor 647 | Invitrogen | A21447 | donkey | NA | 1:1000 |
| anti-mouse IgG Alexa Fluor 488 | Invitrogen | A21202 | donkey | NA | 1:1000 |
| anti-mouse IgG Alexa Fluor 555 | Invitrogen | A31570 | donkey | NA | 1:500 |
| anti-rabbit IgG Alexa Fluor 488 | Invitrogen | A21206 | donkey | NA | 1:1000 |
| anti-rabbit IgG Alexa Fluor 555 | Invitrogen | A31572 | donkey | NA | 1:500 |
| α-SMA | Invitrogen | PA5-18292 | goat | *polyclonal* | 1:100 |
| CD31 | Sigma-Aldrich | P8590 | mouse | WM-59 | 1:200 |
| EpCAM | Cell Signaling Technology | 2929 | mouse | VU1D9 | 1:400 |
| EpCAM | Cell Signaling Technology | 36746 | rabbit | D4K8R | 1:100 |
| GFP | Abcam | ab6556 | rabbit | *polyclonal* | 1:200 |
| HLA-G | Exbio | 11-291-C100 | mouse | MEM-G/1 | 1:100 |
| MMP2 | Cell Signaling Technology | 40994 | rabbit | D4M2N | 1:200 |
